# DINOSim: Zero-Shot Object Detection and Semantic Segmentation on Microscopy Images

**DOI:** 10.1101/2025.03.09.642092

**Authors:** Aitor González-Marfil, Estibaliz Gómez-de-Mariscal, Ignacio Arganda-Carreras

## Abstract

We present DINOSim, a novel method for detecting and segmenting objects in microscopy images without the need for large annotated datasets or additional training. DINOSim builds on the pretrained DINOv2 image encoder, which captures semantic information from images. By comparing the encoder’s features of images patches to those of a user-selected reference, DINOSim generates pseudo-labels that guide object detection and segmentation. Subsequently, a k-nearest neighbors framework is then used to refine predictions across new images. Our experiments show that DINOSim can effectively identify and segment previously unseen objects in diverse microscopy datasets, offering performance comparable to supervised approaches while avoiding the need for costly manual labeling. We also investigate how different choices of user prompts selection and model size affect accuracy and generalization. To make the method widely accessible, we provide an open-source Napari plugin (github.com/AAitorG/napari-DINOSim), enabling researchers to easily apply DINOSim to their own data. Overall, DINOSim offers a fast, flexible and practical solution for bioimage analysis, particularly valuable in resource-constrained settings.

## Introduction

Obtaining labeled data is a costly and time-intensive process in many computer vision applications, posing a significant challenge for training fully supervised models. Few-shot learning (1) has emerged as a solution to this problem, enabling models to effectively learn target tasks using a limited amount of labeled data. Within the scope of few-shot learning, zero-shot learning eliminates the need for labeled examples altogether (2), making it especially valuable in fields where extensive labeled datasets are scarce.

A key enabler of these advancements is the emergence of foundation models (3), which are large-scale deep learning models pre-trained on vast datasets and later adapted for specific tasks. Transformers (4), particularly the Vision Transformer (ViT) (5), have become a preferred architecture for foundation models in both natural language processing (NLP) and computer vision. These models leverage self-supervised learning (SSL) (6, 7), a paradigm that effectively utilizes large amounts of unlabeled data to train models capable of performing various tasks, such as classification and segmentation in computer vision. As a result, foundation models exhibit strong zero-shot and few-shot capabilities (8), demonstrating remarkable generalization to tasks and data distributions outside their original training domains.

A prominent example of a foundation model in computer vision is the Segment Anything Model (SAM) (9). SAM combines an image encoder with a prompting system that uses points, bounding boxes or masks to support object segmentation in images and videos (SAM2 (10)). However, the prompting system of SAM lacks the flexibility of natural language prompts, which could offer more detailed and descriptive segmentation instructions. By default, SAM performs instance segmentation, requiring separate prompts for each object of interest. This limitation becomes cumbersome when dealing with multiple objects in an image or across a set of images. While some efforts have been made to integrate text-based prompts with SAM (11), these approaches often involve prompt engineering, which can introduce additional complexity. Other strategies combine SAM-like models with object detection modules (12). However, these methods typically require fine-tuning the model for specific tasks, thereby reducing the versatility of SAM’s original design and limiting its ability to dynamically select different objects.

Another notable foundation model in computer vision is DINO (self-DIstillation with NO labels) (13), which employs a teacher-student paradigm. In this setup, a “teacher” model guides the learning of a “student” model, enabling iterative learning and progressive improvement. The latest version, DINOv2 (14), introduces a refined training process that produces higher-quality features. These features excel at capturing semantic relationships within natural images, ensuring that similar content is represented consistently, even across different domains or styles. The DINOv2 image encoder has demonstrated outstanding performance in tasks such as segmentation, classification, and depth estimation, achieving these results with minimal fine-tuning or through the addition of task-specific components like classification heads or segmentation decoders (14, 15).

In the life sciences, acquiring and annotating bioimage data is often both costly and time-consuming (16). Bioimages exhibit significant variability due to differences in imaging protocols, including staining techniques, device calibration, and the inherent diversity of biological samples (17). These factors make it challenging –and sometimes impractical– to compile large, comprehensive datasets. For instance, the CEM1.5M dataset (18) represents an important step forward by providing a large and diverse resource for mitochondria segmentation in electron microscopy (EM) images. Nevertheless, many subfields still face severe data scarcity, particularly in the study of rare conditions or pathologies. This lack of high-quality annotated data is a major obstacle in developing foundation models tailored to biological data, since their performance depends on the breadth and quality of the datasets used for training (19).

In NLP, the large language models (LLMs) face similar challenges in specialized fields such as biomedicine or microscopy, where domain-specific training data is scarce (20, 21). Although fine-tuning with domain-specific data can improve performance (22–24), this approach requires considerable computational resources and carefully curated datasets. General-purpose foundation models, such as SAM, may not always outperform specialized models in task-specific benchmarks. However, they offer a key advantage: strong adaptability and generalization across tasks and domains without requiring additional training. This adaptability is especially valuable in bioimage analysis, where acquiring high-quality, annotated datasets is often challenging.

We hypothesize that the embeddings generated by DINOv2 can be harnessed for zero-shot object detection and segmentation of previously unseen objects in bioimages, guided by simple point-based prompts. Specifically, we propose that by leveraging DINOv2’s semantic embedding space, objects similar to a user-selected reference can be detected, even in image modalities and biological domains absent from DINOv2’s original training data. Our method, DINOSim, requires no additional training, fine-tuning, or prompt engineering, making it both practical and accessible.

The main contributions of this work are as follows:

1. We demonstrate that embeddings from a vision foundation model (DINOv2) enable zero-shot object detection and segmentation in six different microscopy datasets, without fine-tuning.
2. We systematically evaluate the trade-offs between model size and detection performance.
3. We analyze the impact of prompt selection and prompt quantity on detection stability and accuracy.
4. We release an open-source, user-friendly Napari plugin (github.com/AAitorG/napari-dinoSim), providing an intuitive graphical interface widely accessible to the life sciences community.

## Related Work

### Zero-shot segmentation and object detection

Zero-shot learning extends few-shot learning by eliminating the need for labeled examples, enabling the segmentation or detection of new categories of objects without annotated training data (1, 2). Similar to few-shot methods, it leverages prior knowledge through transfer learning, but differs by generalizing directly to unseen classes. Broadly, zero-shot approaches can be divided into two main families: projection-based methods and generative methods.

### Projection-based approaches

Projection-based approaches map visual features into a semantic space–often derived from word embeddings or large multimodal models–to establish relationships between seen and unseen classes. Grounding DINO (25) exemplifies this category by which generating bounding boxes from textual prompts. Combined with SAM, Grounded SAM (11) uses these bounding boxes as prompts to refine segmentations for a specified textual prompt. Another notable example is CLIPSeg (26), which builds on CLIP (27) by adding a segmentation decoder to produce masks from textual prompts. CLIPSeg can also accept image-based prompts, where a labeled mask highlights the target object. However, when labeled masks are used, the setting shifts from zero-shot to one-shot learning.

### Generative approaches

Generative approaches synthesize features for unseen categories by learning from labeled data and extrapolating to new classes. Many rely on text or word embeddings, such as Word2Vec (28), to guide feature generation. For example, ZS3Net (29) trains a generator that maps word embeddings into features that represent unseen classes, enabling segmentation or detection without explicit labels.

### Interactive and weakly supervised learning

Interactive learning combines model predictions with iterative user feedback, while weakly supervised learning trains models from incomplete or noisy labels. A cost-effective weak supervision strategy is to use small models, such as random forests (30), trained on predefined image features like Sobel or Gaussian filters (31, 32). Convpaint (33) extends this concept by incorporating deep encoder features (e.g., from VGG (34) or DINOv2) within an interactive framework. Users iteratively refine predictions by adding annotations (e.g., doodles) for each class, progressively improving segmentation quality in a guided manner.

Within this landscape, DINOSim aligns with projection-based methods but distinguishes itself by leveraging DINOv2 embeddings for point-based prompting in a truly zero-shot manner, avoiding additional training, fine-tuning, or complex prompt engineering.

## Methods

DINOSim introduces a zero-shot learning approach that leverages the embeddings from DINOv2, which effectively capture the semantic structure of an image. The minimum input required by DINOSim consists of an image and a point prompt that serves as reference for guiding the segmentation (Figure 1a). The image is first mapped into the embedding space using DINOv2, and the coordinates of the point prompt are projected onto that space to obtain its corresponding vector representation. Given this reference vector, DINOSim computes the distance between each position of the embedding and the reference vector. This distance quantifies the semantic similarity between each pixel and the prompt, enabling the pseudo-labeling of each vector in the embedding space with its corresponding distance value.

**Fig. 1.**
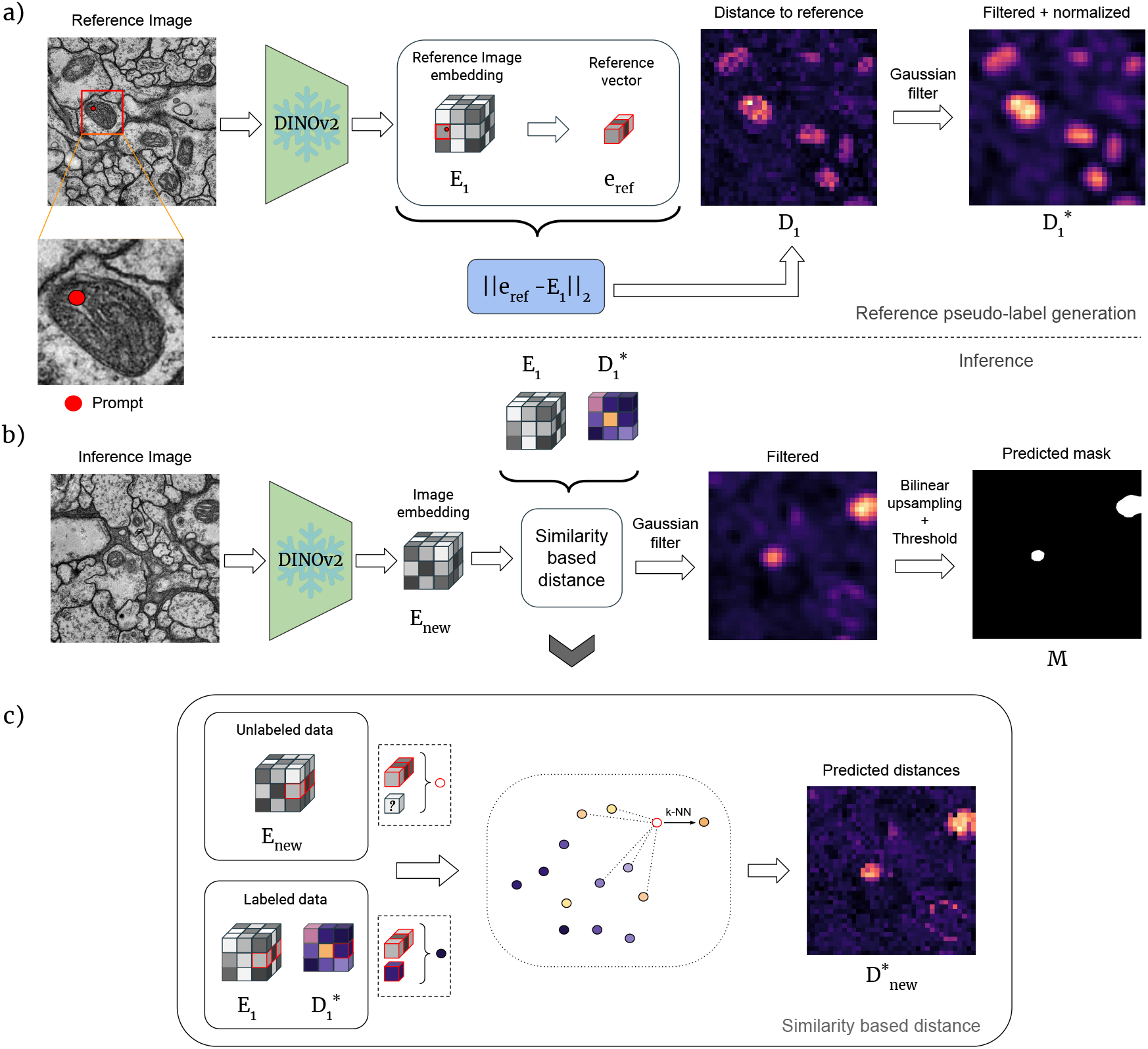
Overview of the DINOSim zero-shot object detection and segmentation method. **(a)** Pseudo-label generation workflow: The image embeddings are extracted using DINOv2. To construct the reference distance map, the Euclidean distance is computed between each patch in the embedding space and the reference vector corresponding to the selected prompt’s coordinates. In the resulting distance map, *D*, lighter areas indicate closer proximity to the reference. The distance map is then smoothed using a Gaussian filter and normalized to obtain *D*^*∗*^. **(b)** Inference workflow: The processed distance map *D*^*∗*^ and the embeddings from the training image set (*{E*_1_, *E*_2_, …, *E*_*T*_ *}*, where *T* = 1 in this case) are used to compute a distance map for a new image via *k*-NN estimation. **(c)** Using *k*-NN, the new distance map is computed as the average of the distance values from the *k*-nearest embedding vectors for each pixel in the embedding space. The resulting distance map is then refined using a Gaussian filter, yielding 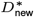. The final segmentation mask, *M*, is obtained by upscaling and thresholding the smoothed distance map.

The pseudo-labeled reference embedding is then used to estimate the similarity values between the embedding of new images and the reference vector. This is achieved through a *k*-nearest neighbor (*k*-NN) approximation which enhances robustness (Figure 1c). Finally, the distance map is filtered, upsampled to match original image resolution, and thresholded to generate the final segmentation mask (Figure 1b). The overall method is illustrated in Figure 1.

### Reference embeddings

Using DINOv2’s image encoder, we extract embeddings from a set of *T* reference images of size of *H* × *W*, each associated with at least one prompt. Since DINOv2 is a ViT, input images are resized to *H*_*p*_ × *W*_*p*_, ensuring compatibility with the model’s patch size. For each reference image *r ∈ {* 1, …, *T}*, its embedding is represented as *E*_*r*_ ∈ ℝ^ℋ×*W×ℱ*^, where *ℋ* and *W* denote the spatial dimensions of the embedding, and *F* represents its feature depth.

### Reference distance maps

For a given reference image *r*, a user-provided set of point-like prompts defines the target elements or regions of interest (ROIs) at spatial coordinates 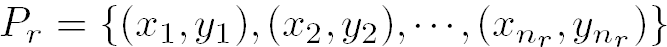. To estimate the feature vector of the prompt *i ∈* [1, …, *n*_*r*_] at location (*x*_*i*_, *y*_*i*_), within the embedding *E*_*r*_, we use bilinear interpolation of the four nearest feature vectors of the point (*x*_*i*_, *y*_*i*_) mapped to the embedding space. This way, the corresponding embedding vectors 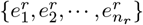 are extracted. The reference vector is computed as the mean of all extracted vectors across all prompts and images:

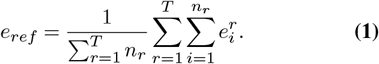

The reference distance map *D*_*r*_ *∈ℝ*^*ℋ*×*W*^ for each image *r* is computed as the Euclidean distance between every vector in *E*_*r*_ and the reference vector *e*_*ref*_ (Figure 1a):

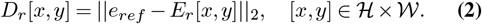

Here, *D*_*r*_ represents the proximity of each pixel to the reference pixels in the embedding space, providing a pseudolabeling map for subsequent segmentation.

### Noise reduction and normalization

Each distance map *D*_*r*_ is smoothed using a Gaussian filter *G*(*·*; *kernel* = 3, *σ* = 1). To mitigate outliers, we apply quantile normalization with a 1% clipping factor, setting the lower and upper clipping thresholds to the 1st and 99th percentiles across all *D*_*r*_ values. The clipped maps,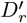, are then normalized to the [0, 1] range using the global minimum and maximum values:

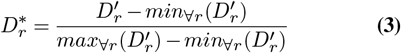

The resulting set of normalized distance maps, 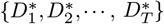, is stored for use in the next step.

#### *k*-NN distance maps

Given a new image and its embedding *E*_*new*_, the embeddings from the reference set *{ E*_1_, *E*_2_, *· · ·, E*_*T*_ *}*are collectively used to determine the *k*-nearest neighbors for each coordinate in *E*_*new*_. The predicted distance value is computed as:

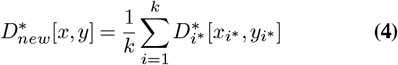

where 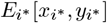 denotes the *i*-th nearest neighbor between *E*_*new*_[*x, y*] among all vectors in *{ E*_1_, *E*_2_, *· · ·, E*_*T*_ *}* (Figure 1c).

#### Post-processing

To refine the predicted distance map 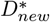, we first apply a Gaussian filter *G*(*·*; *kernel* = 3, *σ* = 1). The smoothed map is then upsampled to the original image resolution (*upsample*(*·*; *H* × *W*)) using bilinear interpolation. The final binary segmentation mask *M* is obtained by thresholding the upsampled distance map with a threshold *τ* (empirically defined):

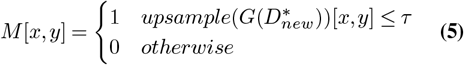

#### Tiling strategy

To efficiently process large images of size *H* ×*W*, we divide them into tiles of size *H*_*t*_ × *W*_*t*_, with the minimum overlap between tiles based in the relation between the total image size and the tile size. Each tile is processed independently through DINOSim up to, but not including, the post-processing step. Before post-processing, the full-size distance map 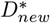 is reconstructed by merging the individual tile predictions. In regions where tiles overlap, pixel values are averaged. Finally, post-processing is applied, and the binary segmentation mask (Eq. 5) is computed at the original image resolution *H* × *W*.

### Benchmarking

#### Benchmarking against fully supervised methods

To assess the feasibility of using DINOv2 image embeddings for microscopy images, we first trained multiple fully supervised models on labeled datasets. Specifically, we implemented: *k*-nearest neighbors (*k*-NN) with *k* = 5, Naive Bayes (NB), a decision tree (DT), a random forest (RF) and a multi-layer perceptron (MLP) with a single hidden layer of 200 neurons. These models were trained to classify embedding vectors extracted from DINOv2 ViT-g. Given an embedding vector (*E*[*x, y*]), the models learned to predict its corresponding class label.

Training labels were generated by downsampling the ground-truth semantic segmentation masks to the embedding resolution (*ℌ* × *W*) using nearest-neighbor interpolation, ensuring that each embedding vector was paired with a label. Final predictions at the original image resolution (*H*× *W*) were obtained using the same post-processing pipeline as in DINOSim.

For comparison with state-of-the-art fully supervised segmentation methods, we trained a U-Net-like model (35) with four levels, ReLU activations, batch normalization (36), 16 initial filters, and batch size of six. The model was trained for 200 epochs using the binary cross-entropy loss function and the AdamW (37) optimizer with a weight decay of 1*e−* 5 and one-cycle learning rate scheduler (38), reaching a maximum learning rate of 1*e −* 3. Unlike DINOv2, our U-Net model operated on 256 × 256 pixel image crops, or on full images when the image size was lower.

To ensure statistical robustness, each supervised model was trained ten times. We report the mean and standard deviation of the accuracy metrics across these runs.

#### Benchmarking against few-shot and zero-shot methods

We next compared DINOSim with state-of-the-art few-shot and zero-shot methods for segmentation and detection. Specifically, we evaluated CLIPSeg (26) and Convpaint (33) for semantic segmentation, and Grounding DINO (25) for object detection. For a fair comparison, bounding boxes were extracted from segmentation masks using the same procedure across all methods. Results are reported in Tables 2 and 3.

**Table 1.**
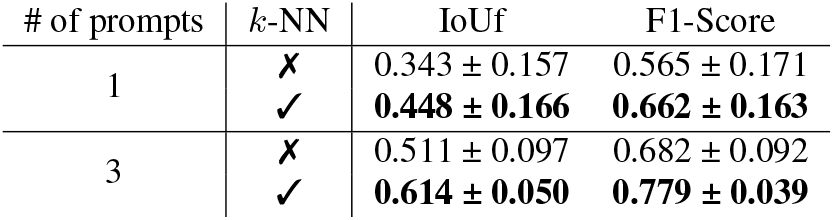
Ablation study on the impact of *k*-NN. Segmentation and detection results on the VNC dataset using different numbers of prompts. Results are reported with and without the *k*-NN step (with *k* = 5) in the inference stage of DINOSim. Embeddings from DINOv2 ViT-g were used. Each value represents the mean ± standard deviation across 100 runs. The best result for each setting is highlighted in bold.

**Table 2.** Semantic segmentation (foreground IoU, IoUf). Comparison across six datasets: VNC, Lucchi++, Staph Aureus (fluorescence “f”, brightfield “bf”), E. coli (brightfield), and NucMM-Mouse. Values are mean *±* s.d. over 10 runs unless noted. DINOSim and CLIPSeg-Img are averaged over 100 runs; CLIPSeg-Txt uses the best textual prompt (single run). A double rule splits the table: (i) fully supervised methods trained on DINOv2 embeddings with vector classifiers (white rows) and U-Net trained on images (light gray), and (ii) few-/zero-shot baselines—few-shot/interactive methods (light gray) and zero-shot methods (dark gray). DINOSim is reported with one (1P) and three (3P) prompts using DINOv2 ViT-g. Bold marks the best score within each section; higher is better.

| Model | IoUf |  |  |  |  |  |
| --- | --- | --- | --- | --- | --- | --- |
|  | VNC | Lucchi++ | Staph Aureus f | Staph Aureus bf | Ecoli bf | NucMM-Mouse |
| DT | 0.546 ± 0.007 | 0.496 ± 0.003 | 0.721 ± 0.002 | 0.682 ± 0.002 | 0.667 ± 0.001 | 0.167 ± 0.009 |
| RF | 0.508 ± 0.002 | 0.404 ± 0.001 | 0.789 ± 0.001 | 0.685 ± 0.001 | 0.674 ± 0.001 | 0.195 ± 0.001 |
| NB | 0.610 | 0.467 | 0.603 | 0.505 | 0.649 | 0.404 |
| k-NN | 0.668 | 0.596 | 0.778 | 0.707 | 0.750 | 0.469 |
| MLP | 0.624 ± 0.003 | 0.586 ± 0.005 | 0.749 ± 0.001 | 0.708 ± 0.002 | 0.720 ± 0.001 | 0.384 ± 0.002 |
| U-Net (35) | **0.839 ± 0.008** | **0.895 ± 0.005** | **0.870 ± 0.003** | **0.722 ± 0.004** | **0.861 ± 0.007** | **0.712 ± 0.011** |
| CLIPSeg-Img (26) | 0.126 ± 0.055 | 0.148 ± 0.045 | 0.061 ± 0.018 | 0.172 ± 0.079 | 0.407 ± 0.091 | 0.032 ± 0.005 |
| ConvPaint (33) | 0.193 ± 0.098 | 0.189 ± 0.063 | 0.223 ± 0.096 | 0.059 ± 0.049 | 0.138 ± 0.115 | 0.033 ± 0.024 |
| CLIPSeg-Txt (26) | 0.059 | 0.067 | 0.377 | 0.339 | 0.108 | 0.002 |
| DINOSim 1P (ours) | 0.448 ± 0.166 | 0.421 ± 0.119 | **0.392 ± 0.077** | 0.336 ± 0.101 | **0.539 ± 0.089** | 0.235 ± 0.096 |
| DINOSim 3P (ours) | **0.614 ± 0.050** | **0.486 ± 0.062** | 0.356 ± 0.056 | **0.341 ± 0.048** | 0.516 ± 0.060 | **0.258 ± 0.073** |

**Table 3.** Object detection (F1-Score). Comparison across six datasets (VNC, Lucchi++, Staph Aureus “f/bf”, E. coli bf, NucMM-Mouse). Values are mean *±* s.d. over 10 runs unless noted. DINOSim and CLIPSeg-Img are averaged over 100 runs; CLIPSeg-Txt and Grounding DINO use the best textual prompt (single run). A double rule separates (i) fully supervised methods on DINOv2 embeddings (white rows) and U-Net on images (light gray) from (ii) few-/zero-shot baselines—few-shot/interactive (light gray) and zero-shot (dark gray). DINOSim results use DINOv2 ViT-g with one (1P) or three (3P) prompts. Detection is computed with an IoU threshold of 0.05 and small boxes filtered as described in the Evaluation section. Bold marks the best score within each section; higher is better.

| Model | F1-Score |  |  |  |  |  |
| --- | --- | --- | --- | --- | --- | --- |
|  | VNC | Lucchi++ | Staph Aureus f | Staph Aureus bf | Ecoli bf | NucMM-Mouse |
| DT | 0.805 ± 0.010 | 0.848 ± 0.005 | 0.940 ± 0.007 | 0.936 ± 0.007 | 0.930 ± 0.003 | 0.360 ± 0.010 |
| RF | 0.773 ± 0.006 | 0.828 ± 0.001 | **0.949 ± 0.003** | 0.933 ± 0.004 | 0.942 ± 0.002 | 0.405 ± 0.002 |
| NB | 0.788 | 0.745 | 0.919 | 0.879 | 0.967 | 0.832 |
| *k*-NN | 0.850 | 0.887 | 0.948 | **0.941** | **0.974** | 0.713 |
| MLP | 0.838 ± 0.004 | 0.888 ± 0.003 | 0.934 ± 0.005 | 0.919 ± 0.009 | 0.964 ± 0.002 | 0.718 ± 0.003 |
| U-Net (35) | **0.909 ± 0.008** | **0.941 ± 0.005** | 0.866 ± 0.006 | 0.927 ± 0.008 | 0.818 ± 0.003 | **0.864 ± 0.006** |
| CLIPSeg-Img (26) | 0.342 ± 0.091 | 0.427 ± 0.116 | 0.173 ± 0.149 | 0.607 ± 0.058 | 0.365 ± 0.026 | 0.011 ± 0.007 |
| Convpaint (33) | 0.366 ± 0.102 | 0.356 ± 0.106 | 0.599 ± 0.229 | 0.179 ± 0.208 | 0.406 ± 0.260 | 0.141 ± 0.103 |
| Grounding DINO (25) | 0.166 | 0.168 | 0 | 0 | 0 | 0 |
| CLIPSeg-Txt (26) | 0.161 | 0.020 | 0.596 | 0.592 | 0.216 | 0.004 |
| DINOSim 1P (ours) | 0.662 ± 0.163 | 0.722 ± 0.120 | **0.779 ± 0.122** | 0.703 ± 0.162 | 0.949 ± 0.088 | 0.672 ± 0.193 |
| DINOSim 3P (ours) | **0.779 ± 0.039** | **0.774 ± 0.042** | 0.738 ± 0.095 | **0.724 ± 0.083** | **0.955 ± 0.045** | **0.752 ± 0.122** |

#### Textual prompts

For text-based methods, we tested several class-related terms such us “mitochondria” and “organelle” in the case of mitochondria segmentation datasets, but found that more descriptive phrases, such as “black object” or “black round object”, yielded better results. These descriptive prompts were therefore used for both Grounding DINO and CLIPSeg. Further details can be found in the Supplementary Material.

#### Image prompts

For CLIPSeg, we also explored image-based prompting strategies, including background blurring or darkening, as suggested in the original paper. The best results were obtained by cropping the object of interest and padding it with a white border. Using this strategy, we computed the mean and standard deviation across 100 randomly selected prompts from the training set. To avoid incomplete or ambiguous cases, we excluded small objects (based on area) and objects touching the image borders. More details can be found in the Supplementary Material.

For segmentation, we applied the fixed threshold of 0.3 recommended in the original CLIPSeg implementation.

#### Scribble prompts

To compare DINOSim with Convpaint under controlled conditions, we generated 10 randomized scribble prompts to minimize human bias. Each prompt was created by:

1. Selecting a random image from the training set,
2. Drawing three random straight lines per class (fore-ground and background) using ground-truth masks.

Background scribbles used a circular brush with 10-pixel radius, while foreground scribbles used a 30-pixel radius. Additional constraints included:

- Scribbles could not cross class boundaries or touch un-connected objects,
- Border regions were excluded to prevent ambiguity,
- Maximum scribble length was set to 300 pixels.

For a fair feature comparison of feature representations, Convpaint was tested using DINOv2 features, which outperformed both VGG and hand-crafted image filters. Specifically, Convpaint employed DINOv2 ViT-S as its feature extractor.

### Experimental Results

#### Datasets

We evaluated our method on six publicly available microscopy datasets for semantic segmentation, covering a range of imaging modalities, organisms, and resolutions.

#### VNC (39)

This dataset focuses on **mitochondria segmentation** in EM images of *Drosophila melanogaster*. It consists of a 4.7× 4.7 ×1 (*µm*)^3^ serial section transmission electron microscopy (ssTEM) volume of the third instar larva ventral nerve cord. Two volumes were acquired, each with 1024 × 1024 × 20 voxels, but only one was annotated. We split the annotated volume along the *x*-axis into two equal parts (512 × 1024 × 20 voxels) for training and testing.

#### Lucchi++ (EPFL Hippocampus) (40, 41)

This dataset is also designed for **mitochondria segmentation**, but in mouse hippocampus tissue. It captures a 5 × 5 × 5 (*µm*)^3^ section of the CA1 region with focused ion beam scanning electron microscopy (FIB-SEM) at an isotropic resolution of 5 nm/voxel. The full dataset contains a 2048 × 1536 × 1065 voxel volume. Mitochondria are labeled in two sub-volumes (165 slices of 1024 × 768 pixels) for training and testing. We used the refined annotation version, **Lucchi++**, curated by neuroscientists and a senior biologist to correct misclassifications and boundary inconsistencies.

#### Staphylococcus aureus (DeepBacs) (42)

Provides paired manually annotated 2D images for the instance segmentation of *S. aureus* bacteria, acquired with a GE HealthCare Deltavision OMX system at 37°C using an Olympus 60 × 1.42 NA oil immersion objective and two PCO Edge 5.5 sCMOS cameras. The dataset contains colocalized fluorescence and brightfield microscopy images of 512 × 512 pixels with a pixel size of 80 nm/px. The fluorescence dataset comprises 7 training and 5 test image pairs of Nile Red-stained live *S. aureus* JE2 cells grown on agarose pads.

#### Escherichia coli (DeepBacs) (43)

Provides paired manually annotated 2D time-lapse images for the segmentation of *E. coli*. Brightfield images were acquired at 1-minute intervals on a Nikon Eclipse Ti-E microscope with a 100× oil immersion objective. Raw images (512 × 512 pixels, 16-bit) were upscaled to 1024 × 1024 pixels (8-bit), resulting in a pixel size of 79 nm/px, to improve segmentation quality. The training and test data consist of 19 and 15 independent frames respectively.

#### NucMM-Mouse (44)

This dataset focuses on **nuclei segmentation** in large-scale 3D micro-CT imaging of the mouse visual cortex. It contains approximately 7,000 nuclei in a 700 × 996 × 968 voxel volume (0.25 mm^3^ physical size). Following the dataset protocol, we used four 192 × 192 × 192 voxel sub-volumes for training and another four for testing.

#### Data preprocessing

Since DINOv2 is a ViT trained with a patch size of 14 × 14 pixels and an image size of 518 × 518 pixels at the end of the training, we adopted a fixed size of 518 × 518 pixels (*H*_*p*_ × *W*_*p*_). For an input image of this size, DINOv2 produces an embedding tensor of 37 × 37 × *ℱ* (*ℌ* ×*W* × *ℱ*), where the feature dimension ℱ varies depending on the model variant: 384 for ViT-S, 768 for ViT-B, 1024 for ViT-L, and 1536 for ViT-g.

Given the variability in resolution and object sizes across datasets, we optimized the patch size (*H*_*t*_ × *W*_*t*_) for each dataset using DINOSim. While exhaustively exploring all patch sizes for every method and dataset was not feasible, we first determined the optimal patch size for each method on the VNC dataset and then extrapolated these values to other datasets. This procedure ensured a fair comparison across methods while keeping experiments tractable.

Finally, all images were preprocessed with 1% quantile normalization to standardize intensity ranges. More details can be found in Supplementary Material.

#### Evaluation

We evaluated the performance of our method on semantic segmentation and object detection tasks.

#### Semantic segmentation

Segmentation results were assessed at the original image resolution (*H* × *W*). Specifically, the upsampled predicted mask *M* from Eq. 5 was compared against the original ground truth annotations.

Performance was quantified using the Intersection over Union (IoU). Rather that computing the mean IoU across all classes–which is typically dominated by the background–we report the foreground IoU (IoUf), which exclusively measures segmentation accuracy for the foreground class.

For 2D datasets, the IoUf was computed for each image and averaged across all images. For volumetric datasets, we computed the IoUf between the predicted and ground truth volumes. When multiple volumes were available, we reported the mean IoUf across them. The volumetric evaluation followed these steps:

- Each 2D slice of the volume was processed independently.
- Predicted masks from all slices were combined to reconstruct the 3D volume.

This volume-based strategy ensures comparability with previous work on EM volumetric datasets, even though DINOSim does not explicitly process the third dimension.

#### Object detection

Bounding boxes were extracted from the predicted mask *M* (Eq. 5) and compared with those derived from the ground truth mask. A predicted box was considered a true positive if its IoU with a ground truth box exceeded a chosen threshold. Detection performance was quantified using the F1-Score, accounting for both precision and recall. Due to the low resolution of the embedding space, predicted bounding boxes often struggle to precisely separate closely positioned objects, leading to low IoU values in crowded regions. To address this limitation, we analyzed the impact of the IoU threshold on true positive classification (Figure 2). Results show that:

**Fig. 2.**
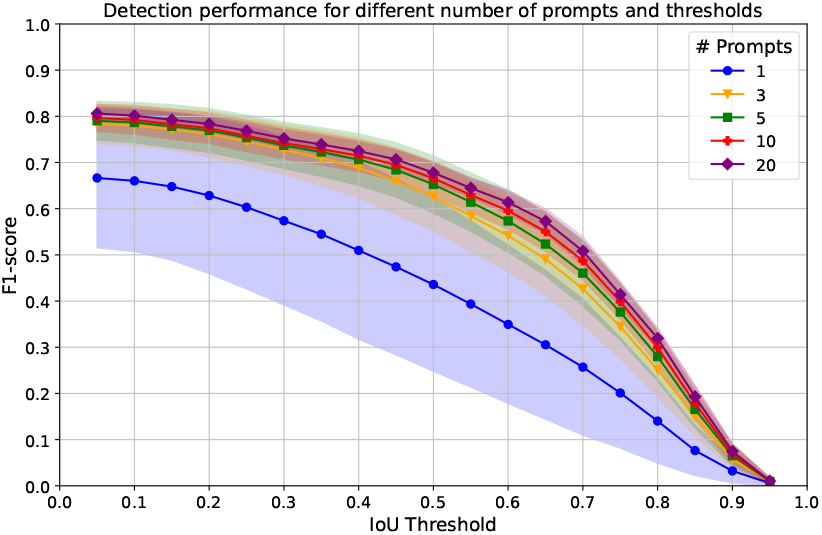
F1-Score across different numbers of prompts and IoU thresholds. The plot shows the average F1-Score across varying IoU thresholds. Different colors denote different numbers prompts. Shaded regions indicate the standard deviation. Results correspond to DINOv2 ViT-g on the VNC dataset, averaged over 100 repetitions. Thresholds range from 0.05 to 0.95 in steps of 0.05.

- Low IoU thresholds reduce the penalization of grouped objects, allowing detections that reasonably enclose multiple targets.
- Overly permissive thresholds could inflate true positives by accepting imprecise, oversized boxes. In such cases, a sharper F1-score decline would be expected as the IoU threshold increases.

Based on these observations, we set the IoU threshold of 0.05 for detection metrics. This permissive threshold accounts for cases where several closely packed objects are grouped into a single predicted box. To further reduce false positives, we excluded detections smaller than approximately 10% of the average object size (more details in Supplementary Material), as these typically correspond to spurious or noise-induced artifacts.

#### Automated prompt selection in DINOSim

To evaluate DINOSim, prompts were selected from the training dataset and used as references to predict masks across the test dataset.

To mimic human prompting behavior while automating the selection process, we designed a sampling strategy that favors prompts located within the object of interest. The sampling relies on a probability map derived from the ground truth segmentation masks, where:

- Background regions and object borders were assigned a probability of zero.
- Probability values increased towards the center of the object, promoting the selection of points well inside the target structures.

This sampling procedure ensured that prompts were consistently positioned in informative, thereby reducing ambiguity and maximizing their relevance for guiding segmentation.

#### Ablation studies

Due to computational constraints, ablation studies were conducted exclusively on EM datasets.

#### Impact of k-NN the inference step

To evaluate the contribution of pseudo-labeling and *k*-NN to DINOSim’s performance, we compared results obtained with and without the *k*-NN step. In the latter case, the computation from Eq. 4 was replaced by directly measuring the Euclidean distance between *E*_*new*_ and the reference vector, as in Eq. 2.

After computing the distance, values were clipped to the range 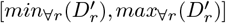 and normalized following Eq. 3, while the post-processing pipeline remained unchanged.

As summarized in Table 1, incorporating *k*-NN with pseudo-labeling clearly improves performance, both in segmentation (IoUf) and object detection (F1-score).

#### Effect of number of prompts and model size

We conducted additional ablation studies to examine how different number of prompts and model sizes contribute to the overall performance of DINOSim in object detection and segmentation. Specifically, we evaluated:

- Varying numbers of prompts (1, 3, 5, 10, and 20).
- Four DINOv2 model variants (ViT-S, ViT-B, ViT-L, and ViT-g), all trained with registers (45).

To avoid possible bias towards a specific dataset, experiments were performed on both VNC and Lucchi++ dataset. Each configuration was repeated 100 times to ensure reliable results.

Object detection results in Figure 3 show that increasing the number of prompts improves the stability of F1-scores across both datasets. However, the effect on overall performance is less consistent.

**Fig. 3.**
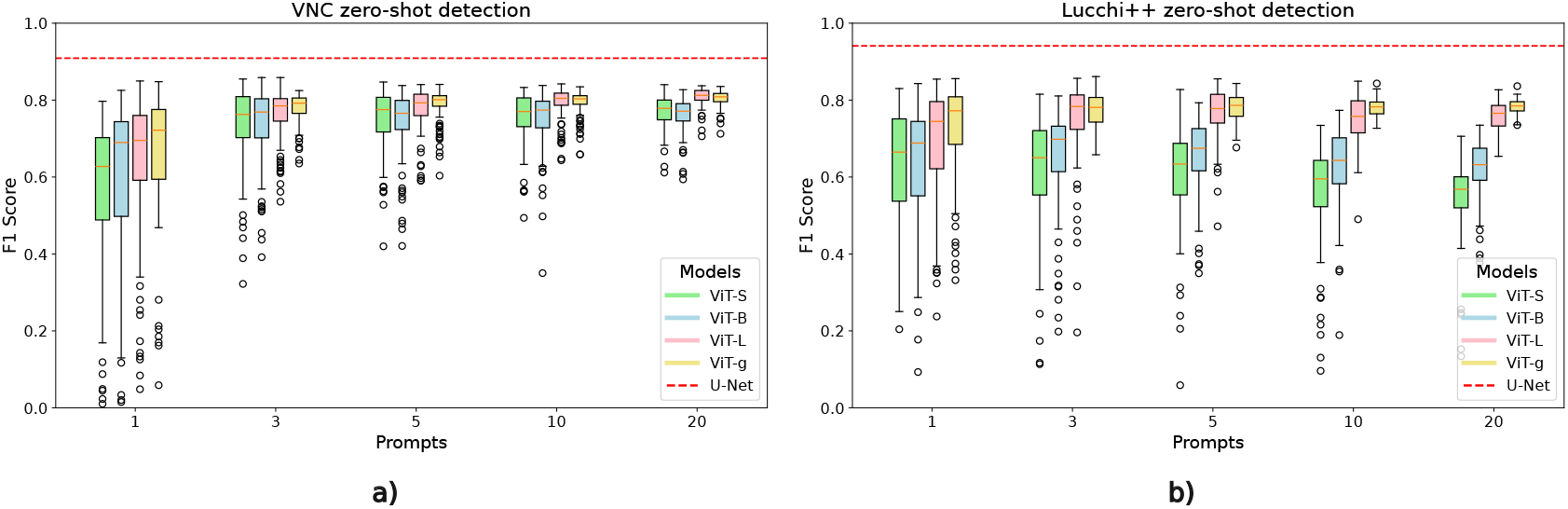
Zero-shot object detection performance with different numbers of prompts and model sizes. F1-scores obtained with DINOSim on the test sets of a) VNC and b) Lucchi++ are shown. The x-axis indicates the number of prompts, while colors represent DINOv2 model sizes. The red line denotes the upper bound, corresponding to the best fully supervised U-Net result. Each box represents 100 runs with randomly selected prompts.

In the VNC dataset, performance generally improves as more prompts are provided. By contrast, in the Lucchi++ dataset, only the two largest models (ViT-L and ViT-g) benefit from additional prompts, while the smaller models exhibit a decline in performance. This suggests that the effect of prompt quantity is both model-dependent and dataset-specific. Nonetheless, results converge quickly, aligning with the observations in Figure 2, where results obtained with three or five prompts are nearly indistinguishable. Regarding model size, larger models demonstrate slightly better and more stable performance.

Since object detection performance is derived from segmentation accuracy, similar trends were observed in both metrics (further details in Supplementary Material).

Although increasing the number of prompts and using larger models may increase runtime, DINOSim leverages GPU acceleration to mitigate computational overhead (execution times are analyzed in Supplementary Material).

For all experiments, the segmentation threshold was set at *τ* = 0.5. However, slight adjustments to *τ* can further improve performance in specific scenarios (see threshold sensitivity analysis in Supplementary Material). Figure 4 illustrates an example zero-shot segmentation result using the default threshold.

**Fig. 4.**
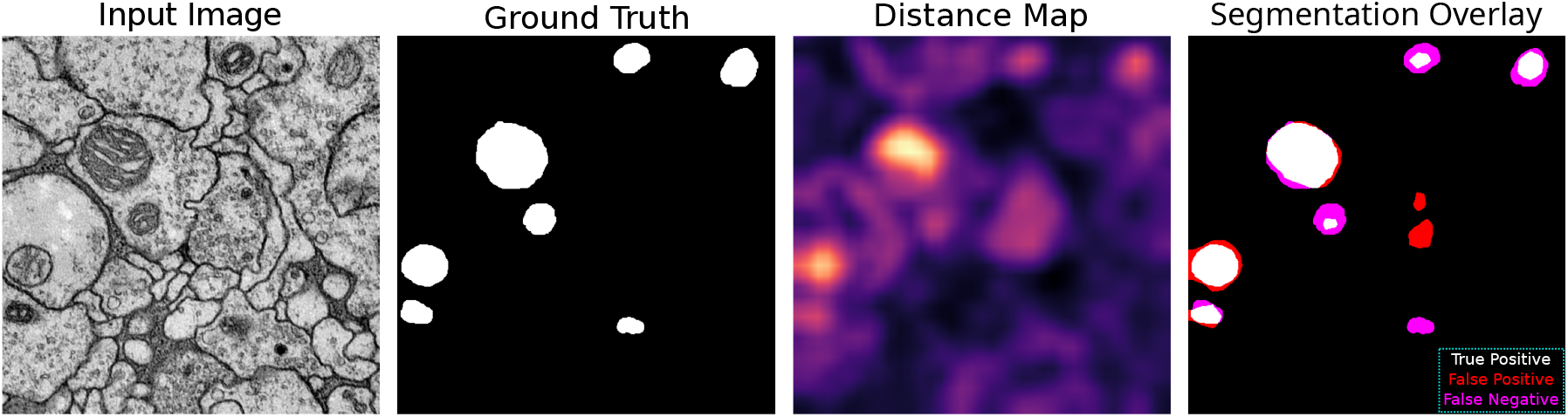
Example of a zero-shot segmentation result with DINOSim. Columns represent: (1) the original image from the VNC test set, (2) the ground truth mask, (3) the upscaled distance map (lighter values indicate higher similarity to the reference), and (4) the predicted mask. In the predicted mask, true positives are shown in white, false positives in red, false negatives in magenta and true negatives in black. This example was generated using DINOv2 ViT-g with 3 randomly selected prompts from the VNC training set.

#### Comparison with supervised and few-shot methods

As shown in Table 2 and Table 3, DINOSim consistently outperforms other few-shot methods in both segmentation and detection, demonstrating robustness and effectiveness in zero-shot learning for biomedical images. With three prompts (DINOSim 3P), performance often approaches or even surpasses that of simpler supervised methods operating on embeddings, highlighting its strong generalization capabilities without requiring extensive labeled data. DINOSim 3P exhibited not only higher mean scores but also lower standard deviation compared to DINOSim 1P, suggesting improved stability and robustness against prompt variability. DINOSim’s relative performance gains were particularly pronounced on challenging datasets like NucMM-Mouse, where it outperformed all few-shot baselines and came within range of supervised methods, underscoring its strength in generalizing to diverse and low-signal biomedical domains.

#### Supervised methods

Focusing on fully supervised models that operate at embedding resolution (top rows of Table 2), *k*-NN generally achieves the strongest results, establishing a reliable upper bound for vector classification methods. This confirms that DINOv2 embeddings effectively cluster similar objects, reflecting their inherent discriminative power. However, operating at embedding resolution significantly constrains segmentation accuracy, leaving all vector classification approaches well behind the U-Net, which sets the state-of-the-art upper bound for pixel-wise segmentation.

Object detection results (Table 3) reveal an interesting contrast: while U-Net excels at segmentation, it is not always the leading method for detection. In particular, on the DeepBacs datasets, several fully supervised embedding-based models achieve detection metrics comparable to, or even higher than, U-Net. Although the margin is small, this highlights DINOv2’s capacity to capture discriminative features useful for object localization across diverse imaging conditions, even without pixel-level supervision.

#### Few-shot and zero-shot methods

Among the few-shot and zero-shot baselines, CLIPSeg with image prompts (CLIPSeg-Img) generally yielded more accurate segmentations than using text prompts (CLIPSeg-Txt), as detailed in Table 2. This indicates the inherent difficulty of describing biomedical structures with purely textual prompts. However, when a suitable textual prompt was found, CLIPSeg-Txt generalized better across modalities, as seen in the fluorescence and brightfield Staph Aureus datasets. This suggests that precise textual descriptions can boost adaptability, though such prompts are not always straightforward to formulate. By contrast, CLIPSeg-Img often oversegmented, producing large regions that encompassed multiple objects, while CLIPSeg-Txt tended to under-segment. These tendencies are illustrated in Figure 5.

**Fig. 5.**
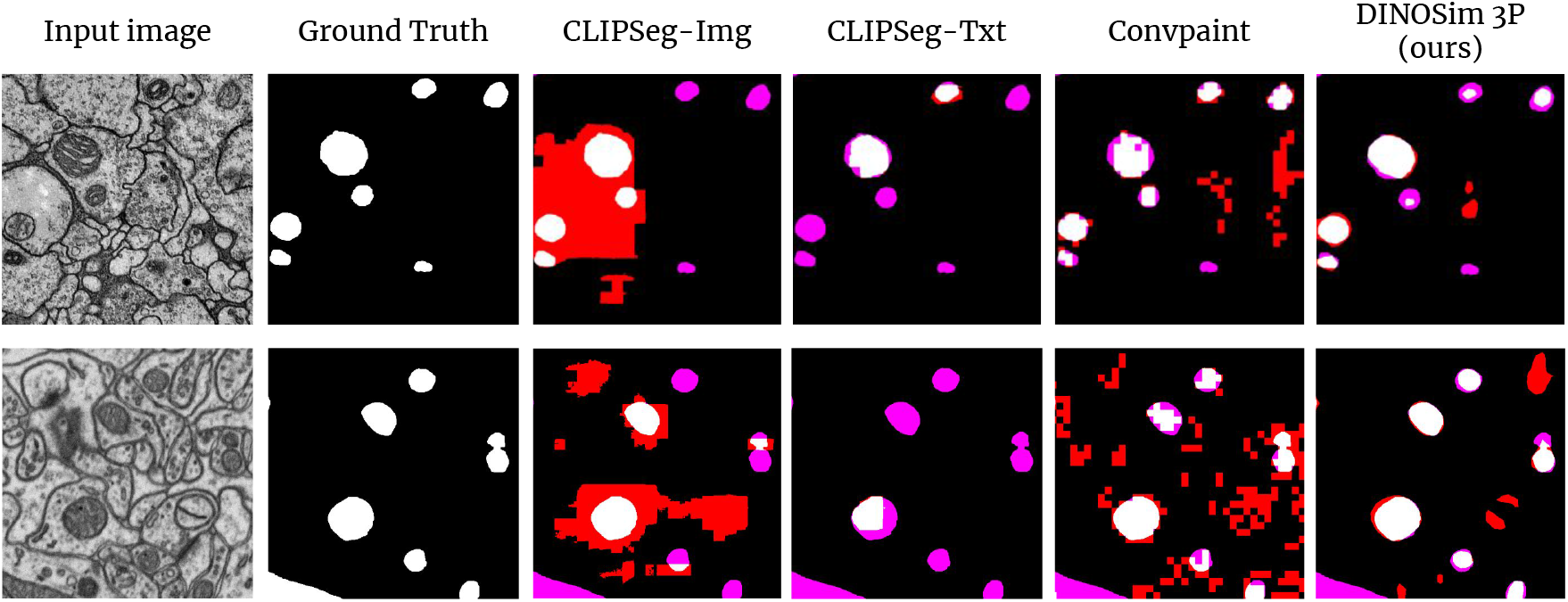
Qualitative comparison of few-shot segmentation methods. From left to right: original image, ground truth mask, predicted masks from CLIPSeg with image prompts, CLIPSeg with textual prompts, Convpaint and DINOSim (ViT-g, 3 prompts). In the predicted masks, true positives are shown in white, false positives in red, false negatives in magenta and true negatives in black. The first row shows an example from the VNC dataset, while the second row shows an example from Lucchi++.

In object detection, Grounding DINO proved largely ineffective, detecting only a small subset of mitochondria across datasets. In contrast, ConvPaint performed better than CLIPSeg across most datasets, likely benefiting from its reliance on DINOv2 embeddings—consistent with the strong performance of DINOv2-based supervised models. However, ConvPaint’s segmentation accuracy was generally comparable to CLIPSeg, and it was more sensitive to imaging shifts, particularly on the Staph Aureus datasets. ConvPaint also exhibited a tendency to oversegment, as seen in Figure 5.

#### Generalization across datasets

We next assessed DINOSim’s robustness to domain shifts by evaluating its performance when prompts (and reference values) were selected from one dataset and tested on another. To reduce variance, we used three references (3P) instead of just one (1P). Results are summarized in Table 4.

**Table 4.** Cross-dataset semantic segmentation and object detection performance. The first column indicates the dataset used for reference selection, while the second column indicates the dataset used for inference. Metrics include segmentation (IoUf) and object detection (F1-Score). Each value is the mean *±* s.d. over 100 runs using DINOv2 ViT-g.

| Dataset |  | DINOSim 3P (ViT-g) |  |
| --- | --- | --- | --- |
| Reference | Inference | IoUf | F1-Score |
| VNC | VNC | 0.614 ± 0.050 | 0.779 ± 0.039 |
| Lucchi++ | VNC | 0.563 ± 0.049 | 0.756 ± 0.043 |
| VNC | Lucchi++ | 0.455 ± 0.056 | 0.769 ± 0.038 |
| Lucchi++ | Lucchi++ | 0.486 ± 0.062 | 0.774 ± 0.042 |

We restricted cross-domain experiments to the EM mitochondria datasets (VNC and Lucchi++) because they share the same biological structure while differing strongly in species and imaging modality. This setup provides a controlled yet challenging test bed for assessing generalization, whereas the remaining datasets target different biological entities, making cross-dataset comparisons less meaningful.

Although both VNC and Lucchi++ focus on mitochondria segmentation, they differ in several key aspects: they originate from different species (fruit fly vs. mouse), they were acquired using different electron microscopy techniques, and they present different structures and contrast distributions. These differences introduce domain shifts, which are well known to degrade the performance of supervised segmentation methods (46). For instance, a U-Net trained on VNC reaches 0.351 *±* 0.101 IoUf on Lucchi++, but when trained on Lucchi++ and tested on VNC, performance collapses to 0.009 *±* 0.010 IoUf.

In contrast, DINOSim maintains stable performance across datasets, demonstrating generalization capabilities in both segmentation and detection tasks. Unlike fully supervised approaches, which are highly sensitive to shifts in imaging conditions and biological context, DINOSim exhibits resilience to dataset variability, exceeding the cross-domain generalization capacity of conventional deep learning approaches.

## Discussion

### Defining the zero-shot boundary

Fair comparison across zero-shot approaches is inherently challenging, as each method relies on distinct prompting strategies. Moreover, the boundary between zero-shot and one-shot learning is often ambiguous.

In image-based prompting, segmentation or recognition of a specific object can be ambiguous when multiple objects are present in the image. A common strategy is to generate object-centric prompts by cropping around the target object. However, this can be interpreted as relaying on detection or segmentation labels, which shifts the method closer to a oneshot setting. From a user’s perspective, the effort required to provide a textual prompt versus selecting a bounding box is nearly equivalent. Furthermore, when prompt engineering is necessary, the need for iterative refinements often makes bounding box prompting a more efficient alternative than textual zero-shot prompting, as it reduces the number of human interactions.

Interactive learning methods complicate this distinction further. While they do not rely on fully annotated datasets, they occupy a gray area between zero-shot and few-shot learning. As the number of iterations grows, these methods demand more training time and annotation effort than standard few-shot approaches. Additionally, they require highly specific human corrections to adjust the model’s predictions.

Paradoxically, interactive methods can become more label-intensive than other few-shot techniques. A further challenge is their tendency to converge toward suboptimal labels, creating uncertainty about when to stop iterating. This uncertainty can lead to excessive relabeling and retraining, ultimately increasing the annotation burden.

### Limitations of DINOSim

DINOSim segments objects based on similarity in the embedding space of a fixed, pre-trained image encoder. This reliance on frozen embeddings limits control over what the model interprets as “similar”. For instance, distinguishing between damaged and non-damaged mitochondria may be difficult, as both categories could be encoded with similar representations. Consequently, they might be segmented together, reducing the methods’ effectiveness for fine-grained classification tasks.

## Conclusions

In this work, we addressed the challenge of scarced labeled data in microscopy by introducing DINOSim, a zero-shot method for object detection and semantic segmentation that leverages DINOv2 embeddings without additional training or labels. Compared to existing few-shot approaches, DINOSim consistently achieved superior performance using only simple point-based prompts.

A key limitation of DINOSim is its reduced effectiveness for pixel-level segmentation tasks when operating directly on embeddings. Inspired by Grounded SAM (11), one possible extension would be to integrate DINOSim with SAM (9, 10) to enhance segmentation precision. While such combination could improve results, it would likely incur higher computational cost and reduced efficiency.

Future directions include investigating multi-resolution strategies or super-resolution in the embedding space (47) to improve fine-grained segmentation, as well as refining object detection by incorporating additional instance or boundary-aware, information.

Despite these challenges, DINOSim offers a simple, efficient, and accessible zero-shot object detection and semantic segmentation solution across diverse imaging conditions and object types. Beyond its immediate usability through an open-source Napari plugin (github.com/AAitorG/napari-DINOSim), DINOSim establishes a foundation for the development of future zero-shot bioimage analysis pipelines, bridging foundation models and practical biomedical applications.

## ACKNOWLEDGEMENTS

This work is partially supported by grant GIU23/022 (to I.A-C.) funded by the University of the Basque Country (UPV/EHU), and grant PID2021-126701OB-I00 (to I.A-C.), funded by the Ministerio de Ciencia, Innovación y Universidades, AEI, MCIN/AEI/10.13039/501100011033, and by “ERDF A way of making Europe”. E.G-de-M. acknowledges the support of the Gulbenkian Foundation (Fundação Calouste Gulbenkian), the European Union through the Horizon Europe program (AI4LIFE project with grant agreement 101057970-AI4LIFE granted to the Optical Cell Biology laboratory at Instituto Gulbenkian de Ciência), the European Molecular Biology Organization (EMBO) Postdoctoral Fellowship (EMBO ALTF 174-2022 to E.G-de-M.) and the Fundação para a Ciência e Tecnologia, Portugal (FCT fellowship 2023.09182.CEECIND to E.G-de-M.). Funded by the European Union. However, the views and opinions expressed are those of the authors only and do not necessarily reflect those of the European Union. Neither the European Union nor the granting authority can be held responsible for them.

## Supplementary Note 1: Analysis of the distance map threshold values

We realized that tuning the threshold used to extract masks out of distance maps, led to better results, but whether moving higher or lower, depends on the data. If we perform a posterior analysis using the ground truth, we can evaluate the performance on segmentation using different thresholds, as we can see in Figure 6. In both datasets with 3 prompts, using 0.5 threshold seems to be the best option. If we evaluate the performance evolution with 1 prompt, that not only the variance is higher, but also the best average IoUf is found with different thresholds, 0.6 in case of VNC and 0.55 in case of Lucchi++. As the best threshold tends to be close to 0.5, we decided to make this the default threshold.

**Fig. 6.**
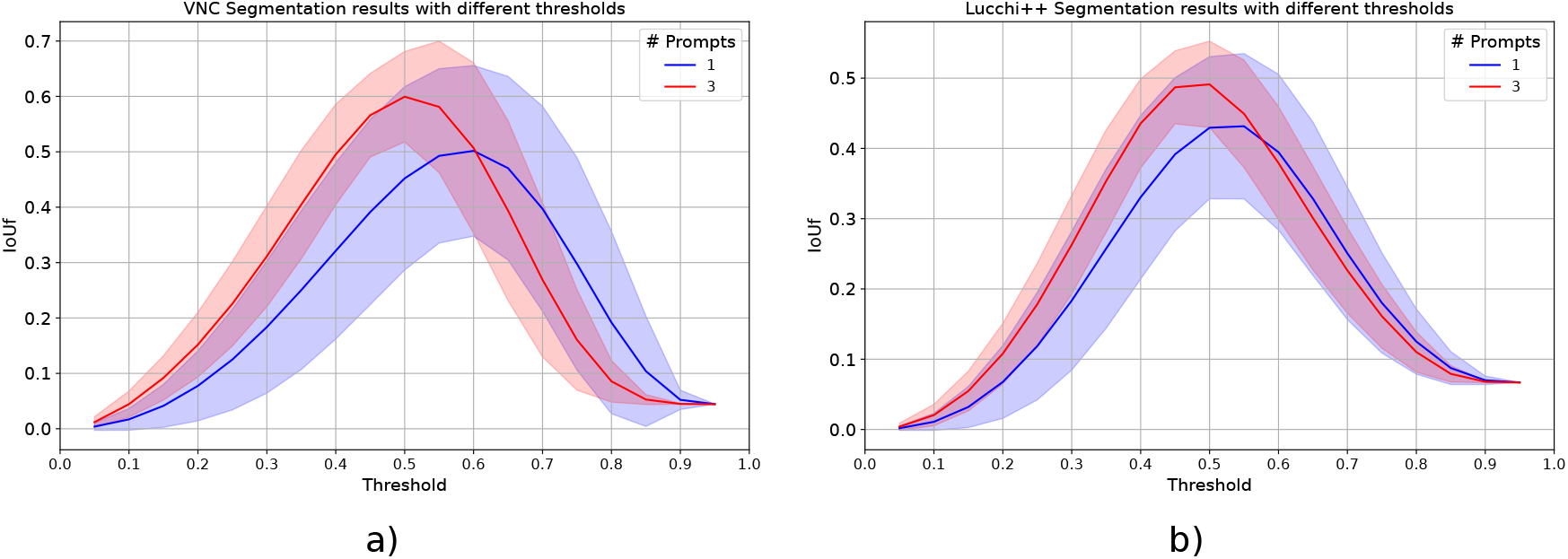
Semantic segmentation performance at different thresholds. The figure shows the IoUf as a function of the threshold values used to extract masks from the predicted distance maps of the VNC test dataset in a) and Lucchi++ test dataset in b), using DINOSim with the ViT-g. The blue line represents the average result from 100 runs using the three prompts, and the shaded blue area shows the standard deviation in these results.

## Supplementary Note 2: Impact of the number of neighbors in k-NN

We evaluated the influence of different number of neighbors in the segmentation and detection performance, in both fully supervised and DINOSim approach, shown in Table 5 respectively. In both scenarios, results are very similar, showing a strong invariance to the number of neighbors. With slightly better results, we selected *k*=5 for the fully supervised method. While in DINOSim, slightly better results appears with more neighbors, this could be counterproductive. As the method works with generated pseudo-labels, may happen to prompted class to belong to a small object, with not many positives. To deal with this unbalance scenario, by default we decided to use a lower amount of neighbors *k*=5.

**Table 5.**
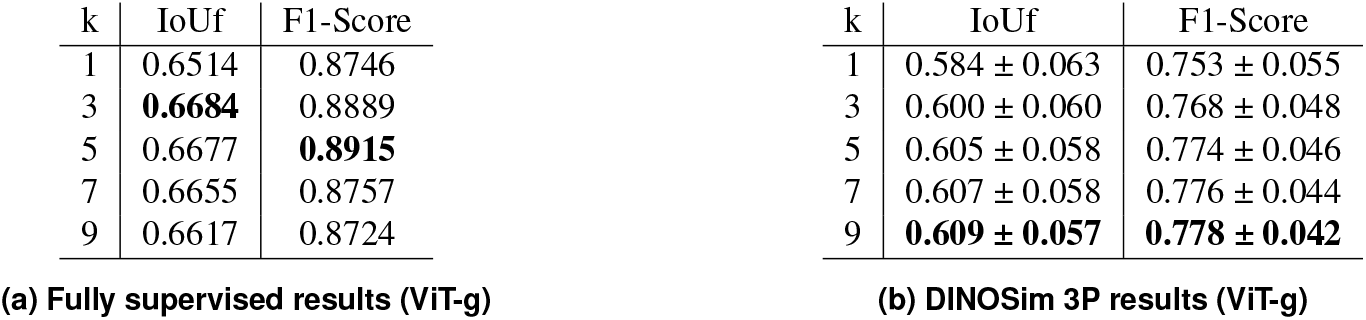
Impact of the number of neighbors in *k*-NN. Semantic segmentation and object detection results using a different number of neighbors in the *k*-NN step for the fully supervised (a) and DINOSim (b) approaches.

## Supplementary Note 3: Impact of the number of prompts and the model size in semantic segmentation results

In this section, variation in segmentation performance for different amount of prompts and different model sizes are evaluated. Regarding the model size, there is a clear tendency of larger models offering better, and more stable segmentation results, see Figure 7. Notably, there is a gap between the two smallest and largest models, the larger the number of prompts the larger the gap. The same conclusions can be seen in detection (see figure 3).

**Fig. 7.**
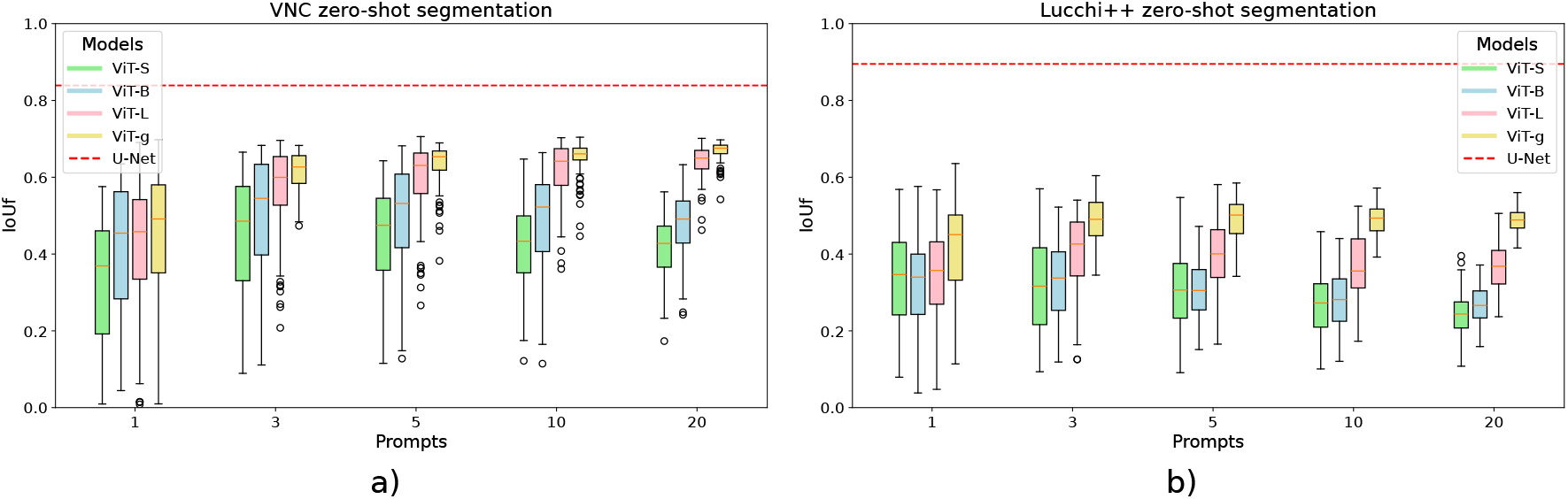
Zero-shot semantic segmentation performance for different prompts. The figure presents the IoUf obtained with DINOSim in the test set of the VNC dataset in a) and Lucchi++ dataset in b), using different number of prompts, while the colors indicate various DINOv2 model sizes. The red line marks the upper bound, showing the best fully supervised result using the U-Net model. Each box represents 100 runs with random prompts.

## Supplementary Note 4: Execution times

All experiments were conducted on a system featuring an Intel Xeon Silver 4210R CPU and an NVIDIA GeForce RTX 3090 GPU with CUDA 12.2. The implementation was built using PyTorch version 2.4.1. Runtime performance details are summarized in Table 6, using the largest model, DINOv2 ViT-g. Since image embeddings are computed only once and can be reused for any number of subsequent prompts, this design ensures that DINOSim operates with improved speed and responsiveness.

**Table 6.**
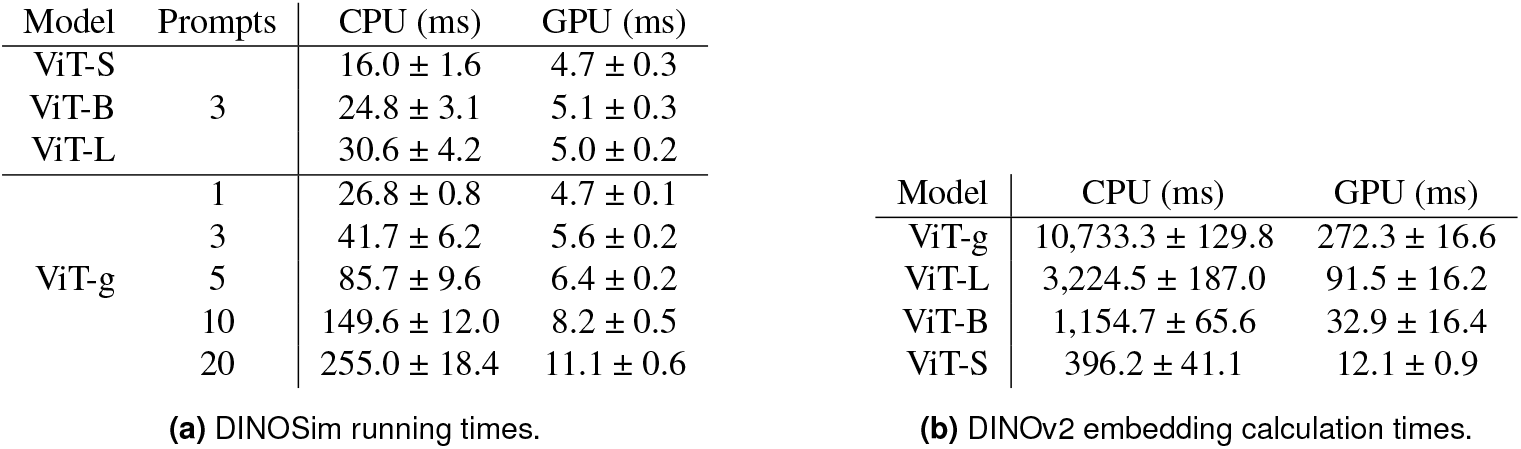
Execution times. a) The table presents, by column: DINOv2 model size, the number of prompts, runtimes on the CPU, and runtimes on the GPU. The reported times reflect the duration required to generate predicted masks using precomputed image embeddings. For CPU runtimes, values were averaged over 400 repetitions, while GPU runtimes were averaged over 4000 repetitions with varying random prompts. b) The table presents, by column: DINOv2 model size, runtimes on the CPU, and runtimes on the GPU. The reported times reflect the duration required to obtain image embeddings using DINOv2. Times were averaged over 40 runs. All times in a) and b) are measured per crop of 518 × 518 pixels.

## Supplementary Note 5: Dataset processing details

Table 7 presents the exact parameter settings used for each dataset.

**Table 7.**
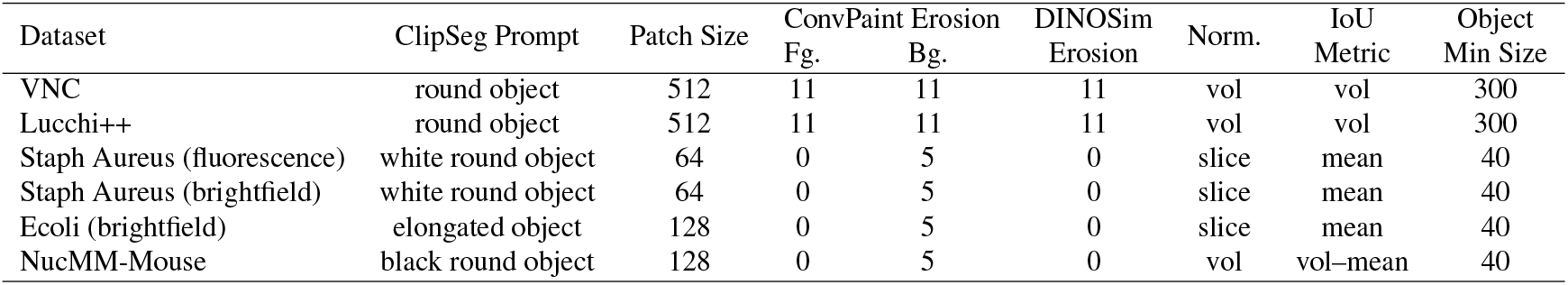
Parameter settings for each dataset.

### Column Descriptions

- **ClipSeg Prompt**: The natural-language phrase used in ClipSeg to obtain the segmentation.
- **Patch Size**: The crop patch size (in pixels) for both *H*_*t*_ and *W*_*t*_.
- **ConvPaint Fg. Erosion**: The disk radius (in pixels) for foreground mask erosion using `skimage.morphology.binary_erosion` prior to prompting.
- **ConvPaint Bg. Erosion**: The disk radius (in pixels) for background mask erosion using `skimage.morphology.binary_erosion` prior to prompting.
- **DINO-SIM Erosion (px)**: The disk/sphere radius for foreground class erosion, used to obtain prompts automatically for DINOSim evaluation. A value of zero indicates no erosion was performed.
- **Normalization**: Indicates whether intensity-scaling (clipping at the 1st percentile and rescaling to [0,1]) was applied per-slice (“slice”) in 2D or per-volume (“vol”) in 3D.
- **IoU Metric**: Specifies whether Intersection-over-Union (IoU) is computed per-slice (with the mean taken across all slices, denoted as “mean”) or on the full 3D volume (denoted as “vol”). For the NucMM-Mouse dataset, due to there are multiple volumes, we measured IoU volume-wise and then computed the mean among all volumes.
- **Object Min Size**: The minimum object area (in pixels) required for an object to be considered valid and not a noise artifact in the detection process.

## Bibliography

1. Simone Antonelli, Danilo Avola, Luigi Cinque, Donato Crisostomi, Gian Luca Foresti, Fabio Galasso, Marco Raoul Marini, Alessio Mecca, and Daniele Pannone. Few-shot object detection: A survey. ACM Comput. Surv., 54(11s), sep 2022. ISSN 0360-0300. doi: 10.1145/3519022.

2. Wenqi Ren, Yang Tang, Qiyu Sun, Chaoqiang Zhao, and Qing-Long Han. Visual semantic segmentation based on few/zero-shot learning: An overview. IEEE/CAA Journal of Automatica Sinica, 11(5):1106–1126, 2024. doi: 10.1109/JAS.2023.123207.

3. Rishi Bommasani, Drew A. Hudson, Ehsan Adeli, Russ Altman, Simran Arora, Sydney von Arx, Michael S. Bernstein, Jeannette Bohg, Antoine Bosselut, Emma Brunskill, Erik Brynjolfsson, Shyamal Buch, Dallas Card, Rodrigo Castellon, Niladri Chatterji, Annie Chen, Kathleen Creel, Jared Quincy Davis, Dora Demszky, Chris Donahue, Moussa Doumbouya, Esin Durmus, Stefano Ermon, John Etchemendy, Kawin Ethayarajh, Li Fei-Fei, Chelsea Finn, Trevor Gale, Lauren Gillespie, Karan Goel, Noah Goodman, Shelby Grossman, Neel Guha, Tatsunori Hashimoto, Peter Henderson, John Hewitt, Daniel E. Ho, Jenny Hong, Kyle Hsu, Jing Huang, Thomas Icard, Saahil Jain, Dan Jurafsky, Pratyusha Kalluri, Siddharth Karamcheti, Geoff Keeling, Fereshte Khani, Omar Khattab, Pang Wei Koh, Mark Krass, Ranjay Krishna, Rohith Kuditipudi, Ananya Kumar, Faisal Ladhak, Mina Lee, Tony Lee, Jure Leskovec, Isabelle Levent, Xiang Lisa Li, Xuechen Li, Tengyu Ma, Ali Malik, Christopher D. Manning, Suvir Mirchandani, Eric Mitchell, Zanele Munyikwa, Suraj Nair, Avanika Narayan, Deepak Narayanan, Ben Newman, Allen Nie, Juan Carlos Niebles, Hamed Nilforoshan, Julian Nyarko, Giray Ogut, Laurel Orr, Isabel Papadimitriou, Joon Sung Park, Chris Piech, Eva Portelance, Christopher Potts, Aditi Raghunathan, Rob Reich, Hongyu Ren, Frieda Rong, Yusuf Roohani, Camilo Ruiz, Jack Ryan, Christopher Ré, Dorsa Sadigh, Shiori Sagawa, Keshav Santhanam, Andy Shih, Krishnan Srinivasan, Alex Tamkin, Rohan Taori, Armin W. Thomas, Florian Tramèr, Rose E. Wang, William Wang, Bohan Wu, Jiajun Wu, Yuhuai Wu, Sang Michael Xie, Michihiro Yasunaga, Jiaxuan You, Matei Zaharia, Michael Zhang, Tianyi Zhang, Xikun Zhang, Yuhui Zhang, Lucia Zheng, Kaitlyn Zhou, and Percy Liang. On the opportunities and risks of foundation models, 2022.

4. Ashish Vaswani, Noam Shazeer, Niki Parmar, Jakob Uszkoreit, Llion Jones, Aidan N Gomez, Łukasz Kaiser, and Illia Polosukhin. Attention is all you need. In I. Guyon, U. Von Luxburg, S. Bengio, H. Wallach, R. Fergus, S. Vishwanathan, and R. Garnett, editors, Advances in Neural Information Processing Systems, volume 30. Curran Associates, Inc., 2017.

5. Alexey Dosovitskiy, Lucas Beyer, Alexander Kolesnikov, Dirk Weissenborn, Xiaohua Zhai, Thomas Unterthiner, Mostafa Dehghani, Matthias Minderer, Georg Heigold, Sylvain Gelly, Jakob Uszkoreit, and Neil Houlsby. An image is worth 16×16 words: Transformers for image recognition at scale, 2021.

6. Randall Balestriero, Mark Ibrahim, Vlad Sobal, Ari Morcos, Shashank Shekhar, Tom Goldstein, Florian Bordes, Adrien Bardes, Gregoire Mialon, Yuandong Tian, Avi Schwarzschild, Andrew Gordon Wilson, Jonas Geiping, Quentin Garrido, Pierre Fernandez, Amir Bar, Hamed Pirsiavash, Yann LeCun, and Micah Goldblum. A cookbook of self-supervised learning, 2023.

7. Kaiming He, Xinlei Chen, Saining Xie, Yanghao Li, Piotr Dollár, and Ross Girshick. Masked autoencoders are scalable vision learners. In Proceedings of the IEEE/CVF Conference on Computer Vision and Pattern Recognition (CVPR), pages 16000–16009, June 2022.

8. Tom Brown, Benjamin Mann, Nick Ryder, Melanie Subbiah, Jared D Kaplan, Prafulla Dhariwal, Arvind Neelakantan, Pranav Shyam, Girish Sastry, Amanda Askell, Sandhini Agarwal, Ariel Herbert-Voss, Gretchen Krueger, Tom Henighan, Rewon Child, Aditya Ramesh, Daniel Ziegler, Jeffrey Wu, Clemens Winter, Chris Hesse, Mark Chen, Eric Sigler, Mateusz Litwin, Scott Gray, Benjamin Chess, Jack Clark, Christopher Berner, Sam McCandlish, Alec Radford, Ilya Sutskever, and Dario Amodei. Language models are few-shot learners. In H. Larochelle, M. Ranzato, R. Hadsell, M.F. Balcan, and H. Lin, editors, Advances in Neural Information Processing Systems, volume 33, pages 1877–1901. Curran Associates, Inc., 2020.

9. Alexander Kirillov, Eric Mintun, Nikhila Ravi, Hanzi Mao, Chloe Rolland, Laura Gustafson, Tete Xiao, Spencer Whitehead, Alexander C. Berg, Wan-Yen Lo, Piotr Dollar, and Ross Girshick. Segment anything. In Proceedings of the IEEE/CVF International Conference on Computer Vision (ICCV), pages 4015–4026, October 2023.

10. Nikhila Ravi, Valentin Gabeur, Yuan-Ting Hu, Ronghang Hu, Chaitanya Ryali, Tengyu Ma, Haitham Khedr, Roman Rädle, Chloe Rolland, Laura Gustafson, Eric Mintun, Junting Pan, Kalyan Vasudev Alwala, Nicolas Carion, Chao-Yuan Wu, Ross Girshick, Piotr Dollár, and Christoph Feichtenhofer. Sam 2: Segment anything in images and videos, 2024.

11. Tianhe Ren, Shilong Liu, Ailing Zeng, Jing Lin, Kunchang Li, He Cao, Jiayu Chen, Xinyu Huang, Yukang Chen, Feng Yan, Zhaoyang Zeng, Hao Zhang, Feng Li, Jie Yang, Hongyang Li, Qing Jiang, and Lei Zhang. Grounded sam: Assembling open-world models for diverse visual tasks, 2024.

12. Uriah Israel, Markus Marks, Rohit Dilip, Qilin Li, Morgan Schwartz, Elora Pradhan, Edward Pao, Shenyi Li, Alexander Pearson-Goulart, Pietro Perona, Georgia Gkioxari, Ross Barnowski, Yisong Yue, and David Van Valen. A foundation model for cell segmentation, 2023.

13. Mathilde Caron, Hugo Touvron, Ishan Misra, Hervé Jégou, Julien Mairal, Piotr Bojanowski, and Armand Joulin. Emerging properties in self-supervised vision transformers. In Proceedings of the IEEE/CVF International Conference on Computer Vision (ICCV), pages 9650–9660, October 2021.

14. Maxime Oquab, Timothée Darcet, Théo Moutakanni, Huy Vo, Marc Szafraniec, Vasil Khalidov, Pierre Fernandez, Daniel Haziza, Francisco Massa, Alaaeldin El-Nouby, Mahmoud Assran, Nicolas Ballas, Wojciech Galuba, Russell Howes, Po-Yao Huang, Shang-Wen Li, Ishan Misra, Michael Rabbat, Vasu Sharma, Gabriel Synnaeve, Hu Xu, Hervé Jegou, Julien Mairal, Patrick Labatut, Armand Joulin, and Piotr Bojanowski. Dinov2: Learning robust visual features without supervision, 2024.

15. Joana Palés Huix, Adithya Raju Ganeshan, Johan Fredin Haslum, Magnus Söderberg, Christos Matsoukas, and Kevin Smith. Are natural domain foundation models useful for medical image classification? In Proceedings of the IEEE/CVF Winter Conference on Applications of Computer Vision (WACV), pages 7634–7643, January 2024.

16. Chongyu Qu, Tiezheng Zhang, Hualin Qiao, Jie Liu, Yucheng Tang, Alan Yuille, and Zongwei Zhou. Abdomenatlas-8k: Annotating 8,000 ct volumes for multi-organ segmentation in three weeks, 2023.

17. Melissa Linkert, Curtis T. Rueden, Chris Allan, Jean-Marie Burel, Will Moore, Andrew Patterson, Brian Loranger, Josh Moore, Carlos Neves, Donald MacDonald, Aleksandra Tarkowska, Caitlin Sticco, Emma Hill, Mike Rossner, Kevin W. Eliceiri, and Jason R. Swedlow. Metadata matters: access to image data in the real world. Journal of Cell Biology, 189 (5):777–782, 05 2010. ISSN 0021-9525. doi: 10.1083/jcb.201004104.

18. Ryan Conrad and Kedar Narayan. Instance segmentation of mitochondria in electron microscopy images with a generalist deep learning model trained on a diverse dataset. Cell Systems, 14(1):58–71.e5, Jan 2023. ISSN 2405-4712. doi: 10.1016/j.cels.2022.12.006.

19. Jiaxing Huang, Kai Jiang, Jingyi Zhang, Han Qiu, Lewei Lu, Shijian Lu, and Eric Xing. Learning to prompt segment anything models, 2024.

20. Zabir Al Nazi and Wei Peng. Large language models in healthcare and medical domain: A review. Informatics, 11(3), 2024. ISSN 2227-9709. doi: 10.3390/informatics11030057.

21. Md Adnan Arefeen, Biplob Debnath, and Srimat Chakradhar. Leancontext: Cost-efficient domain-specific question answering using llms. Natural Language Processing Journal, 7: 100065, 2024. ISSN 2949-7191. doi: 10.1016/j.nlp.2024.100065.

22. Karan Singhal, Tao Tu, Juraj Gottweis, Rory Sayres, Ellery Wulczyn, L. Hou, Kevin Clark, Stephen Pfohl, Heather Cole-Lewis, Darlene Neal, Mike Schaekermann, Amy Wang, Mohamed Amin, Sami Lachgar, Philip Mansfield, Sushant Prakash, Bradley Green, Ewa Dominowska, Blaise Aguera y Arcas, Nenad Tomasev, Yun Liu, Renee Wong, Christopher Semturs, S. Sara Mahdavi, Joelle Barral, Dale Webster, Greg S. Corrado, Yossi Matias, Shekoofeh Azizi, Alan Karthikesalingam, and Vivek Natarajan. Towards expert-level medical question answering with large language models, 2023.

23. Jun Ma, Yuting He, Feifei Li, Lin Han, Chenyu You, and Bo Wang. Segment anything in medical images. Nature Communications, 15(1):654, Jan 2024. ISSN 2041-1723. doi: 10.1038/s41467-024-44824-z.

24. Anwai Archit, Sushmita Nair, Nabeel Khalid, Paul Hilt, Vikas Rajashekar, Marei Freitag, Sagnik Gupta, Andreas Dengel, Sheraz Ahmed, and Constantin Pape. Segment anything for microscopy. bioRxiv, 2023. doi: 10.1101/2023.08.21.554208.

25. Shilong Liu, Zhaoyang Zeng, Tianhe Ren, Feng Li, Hao Zhang, Jie Yang, Qing Jiang, Chunyuan Li, Jianwei Yang, Hang Su, Jun Zhu, and Lei Zhang. Grounding dino: Marrying dino with grounded pre-training for open-set object detection, 2024.

26. Timo Lüddecke and Alexander Ecker. Image segmentation using text and image prompts. In Proceedings of the IEEE/CVF Conference on Computer Vision and Pattern Recognition (CVPR), pages 7086–7096, June 2022.

27. Alec Radford, Jong Wook Kim, Chris Hallacy, Aditya Ramesh, Gabriel Goh, Sandhini Agarwal, Girish Sastry, Amanda Askell, Pamela Mishkin, Jack Clark, Gretchen Krueger, and Ilya Sutskever. Learning transferable visual models from natural language supervision. In Marina Meila and Tong Zhang, editors, Proceedings of the 38th International Conference on Machine Learning, volume 139 of Proceedings of Machine Learning Research, pages 8748–8763. PMLR, 18–24 Jul 2021.

28. Xin Rong. word2vec parameter learning explained, 2016.

29. Maxime Bucher, Tuan-Hung VU, Matthieu Cord, and Patrick Pérez. Zero-shot semantic segmentation. In H. Wallach, H. Larochelle, A. Beygelzimer, F. d’Alché-Buc, E. Fox, and R. Garnett, editors, Advances in Neural Information Processing Systems, volume 32. Curran Associates, Inc., 2019.

30. Tin Kam Ho. Random decision forests. In Proceedings of 3rd International Conference on Document Analysis and Recognition, volume 1, pages 278–282 vol.1, 1995. doi: 10.1109/ICDAR.1995.598994.

31. Ignacio Arganda-Carreras, Verena Kaynig, Curtis Rueden, Kevin W Eliceiri, Johannes Schindelin, Albert Cardona, and H Sebastian Seung. Trainable Weka Segmentation: a machine learning tool for microscopy pixel classification. Bioinformatics, 33(15):2424–2426, 03 2017. ISSN 1367-4803. doi: 10.1093/bioinformatics/btx180.

32. Stuart Berg, Dominik Kutra, Thorben Kroeger, Christoph N Straehle, Bernhard X Kausler, Carsten Haubold, Martin Schiegg, Janez Ales, Thorsten Beier, Markus Rudy, Kemal Eren, Jaime I Cervantes, Buote Xu, Fynn Beuttenmueller, Adrian Wolny, Chong Zhang, Ullrich Koethe, Fred A Hamprecht, and Anna Kreshuk. ilastik: interactive machine learning for (bio)image analysis. Nature Methods, 16(12):1226–1232, December 2019.

33. Lucien Hinderling, Guillaume Witz, Roman Schwob, Ana Stojiljković, Maciej Dobrzyński, Mykhailo Vladymyrov, Joël Frei, Benjamin Grädel, Agne Frismantiene, and Olivier Pertz. Convpaint - interactive pixel classification using pretrained neural networks. bioRxiv, 2024. doi: 10.1101/2024.09.12.610926.

34. Karen Simonyan and Andrew Zisserman. Very deep convolutional networks for large-scale image recognition, 2015.

35. Daniel Franco-Barranco, Arrate Muñoz-Barrutia, and Ignacio Arganda-Carreras. Stable deep neural network architectures for mitochondria segmentation on electron microscopy volumes. Neuroinformatics, 20(2):437–450, Apr 2022. ISSN 1559-0089. doi: 10.1007/s12021-021-09556-1.

36. Sergey Ioffe. Batch normalization: Accelerating deep network training by reducing internal covariate shift. arXiv preprint 1502.03167, 2015.

37. Ilya Loshchilov and Frank Hutter. Decoupled weight decay regularization, 2019.

38. Leslie N. Smith and Nicholay Topin. Super-convergence: Very fast training of neural networks using large learning rates, 2018.

39. Stephan Gerhard, Jan Funke, Julien Martel, Albert Cardona, and Richard Fetter. Segmented anisotropic sstem dataset of neural tissue. figshare, pages 0–0, 2013.

40. Aurélien Lucchi, Kevin Smith, Radhakrishna Achanta, Graham Knott, and Pascal Fua. Supervoxel-based segmentation of mitochondria in em image stacks with learned shape features. IEEE Transactions on Medical Imaging, 31(2):474–486, 2012. doi: 10.1109/TMI.2011.2171705.

41. Vincent Casser, Kai Kang, Hanspeter Pfister, and Daniel Haehn. Fast mitochondria detection for connectomics. In Tal Arbel, Ismail Ben Ayed, Marleen de Bruijne, Maxime Descoteaux, Herve Lombaert, and Christopher Pal, editors, Proceedings of the Third Conference on Medical Imaging with Deep Learning, volume 121 of Proceedings of Machine Learning Research, pages 111–120. PMLR, 06–08 Jul 2020.

42. Pedro Matos Pereira and Mariana Pinho. Deepbacs – staphylococcus aureus widefield segmentation dataset, October 2021.

43. Christoph Spahn and Mike Heilemann. Deepbacs – escherichia coli bright field segmentation dataset, October 2021.

44. Zudi Lin, Donglai Wei, Mariela D Petkova, Yuelong Wu, Zergham Ahmed, Silin Zou, Nils Wendt, Jonathan Boulanger-Weill, Xueying Wang, Nagaraju Dhanyasi, et al. Nucmm dataset: 3d neuronal nuclei instance segmentation at sub-cubic millimeter scale. In International Conference on Medical Image Computing and Computer-Assisted Intervention, pages 164–174. Springer, 2021.

45. Timothée Darcet, Maxime Oquab, Julien Mairal, and Piotr Bojanowski. Vision transformers need registers, 2024.

46. Daniel Franco-Barranco, Julio Pastor-Tronch, Aitor González-Marfil, Arrate Muñoz-Barrutia, and Ignacio Arganda-Carreras. Deep learning based domain adaptation for mitochondria segmentation on em volumes. Computer Methods and Programs in Biomedicine, 222: 106949, 2022. ISSN 0169-2607. doi: 10.1016/j.cmpb.2022.106949.

47. Stephanie Fu, Mark Hamilton, Laura Brandt, Axel Feldman, Zhoutong Zhang, and William T. Freeman. Featup: A model-agnostic framework for features at any resolution, 2024.

